# A global survey of System Biology-based predictions of gene-rare disease associations to enhance new diagnoses

**DOI:** 10.64898/2026.02.09.701767

**Authors:** Yolanda Benítez, Graciela Uría-Regojo, Pablo Mínguez

## Abstract

In rare disease diagnosis, described genotype-phenotype associations are evaluated first. In the absence of strong evidence, WES and WGS provide hundred to million other genetic variants, most poorly annotated, that need to be prioritized. While several *in silico* approaches leverage existing gene-disease knowledge to predict novel associations, doing so in isolation can hide how different genes are represented across other predictions. We hypothesize that a global perspective, accounting for differences in the knowledge accumulated in the gene collections, can refine predictions.

Using a network-based algorithm, we explored functional neighborhoods of known disease-associated genes to predict novel candidates for over 200 rare diseases. A global analysis of gene and protein family behavior across predictions identified genes and functions broadly associated with multiple conditions, 192 genes linked to a single disease and 251 genes functionally associated with specific classes of rare diseases. These findings are integrated into a gene-disease specificity score, aimed at enhancing variant prioritization and guiding geneticists in advancing candidate genes toward functional validation.

## INTRODUCTION

Drawing on methodology developed for protein function prediction [1], computational algorithms can aid in uncovering novel gene-disease associations [2]. A known bias of these methods is derived from the different grade of knowledge accumulated over the human gene collection in databases and articles [3]. Nevertheless, and while not conclusive, their predictions can guide experimental approaches and narrow down the set of candidate targets. This is particularly relevant in rare genetic diseases (RDs) where patients present rare, sometimes unique, genotype-phenotype associations causing disorders [4]. With over 7,000 RDs described most of them monogenic, and on many occasions with small disperse cohorts, the discovery of a new gene involved in this type of traits represents an important challenge. Moreover, the contribution of most genomic variants in disease phenotypes is still poorly understood [5][6], with a large proportion of variants classified as variants of uncertain significance (VUS). As a result, a high degree of uncertainty persists during the genetic diagnosis of RDs once all the known associations have been discarded.

Whole exome (WES) and whole genome (WGS) sequencing provide the possibility to explore genotype-phenotype associations beyond the current knowledge. They are increasingly being proposed, or already established, as first tier tests in the protocols for genetic diagnosis [7] replacing targeted gene panels as the clinical exome (CES) [8]. This shift is driven not only by the decrease in sequencing costs but also by the broader benefits these methods offer [9]. These benefits include the possibility of diagnosing additional present or future disorders, and the integration of preventive information into clinical records as polygenic risk scores [10] or pharmacogenetic insights [11]. Other advantages include contributing to population genetics studies [12][13] and enhancing variant prioritization through cohort-based analyses [14]. Nonetheless, their diagnostic yield is still mostly limited to the regular updates of knowledge in the annotation resources as well as to improved bioinformatics pipelines [15][16].

To strengthen the diagnostic impact of implementing WES and WGS in RDs, these approaches must be complemented with strategies that aid in the prioritization of variants beyond those in known disease associated genes. The special features of RDs and the urgency to provide an answer to long-term delayed diagnosis, require finely tuned methods. Several areas of active research aim to reduce the uncertainty inherent in WES and WGS analyses. Notable examples include the characterization of splice sites [17][18], the annotation of non-coding variants [19], the reclassification of VUS [20], or the prioritization of candidate genes [21][22]. In this context, our laboratory has developed GLOWgenes, a Systems Biology inspired computational algorithm that performs candidate gene prediction optimized for RDs as benchmarked together with other similar tools [23]. GLOWgenes is currently employed to prioritize variants outside of known associated genes in the reanalysis of undiagnosed RD cases integrated within our DNASeq pipeline [24]. Here, we present a global survey of gene-disease association predictions for over 200 RDs using GLOWgenes. This provides not only a good community resource to prioritize new candidate genes in WES and WGS analyses, but also enabled a global analysis to uncover general trends in gene–disease associations that cannot be observed through isolated analyses. We used this information to describe gene and protein family behavior across predictions in RDs and to identify genes that are uniquely associated with a RD, to a family of RDs or broadly distributed. All this knowledge is integrated into a comprehensive score designed to enhance variant prioritization in RD diagnosis, ultimately supporting geneticists in upgrading candidate genes for further functional characterization.Here, we present a global survey of gene-disease association predictions for over 200 RDs using GLOWgenes. This provides not only a good community resource to prioritize new candidate genes in WES and WGS analyses, but also enabled a global analysis to uncover general trends in gene–disease associations that cannot be observed through isolated analyses. We used this information to describe gene and protein family behavior across predictions in RDs and to identify genes that are uniquely associated with a RD, to a family of RDs or broadly distributed. All this knowledge is integrated into a comprehensive score designed to enhance variant prioritization in RD diagnosis, ultimately supporting geneticists in upgrading candidate genes for further functional characterization.

## MATERIALS AND METHODS

### Known gene-disease associations

We downloaded gene sets associated with RDs from the Genomics England PanelApp resource [22]. These sets (also known as gene panels) describe the current knowledge of gene-RD associations and are used as diagnostic tools worldwide. Within the PanelApp panels, genes are classified according to their association to the disease using a color code. We selected green and amber genes as those having strong and some evidence of association to the disease respectively. Candidate (red) genes were excluded. We also excluded super panels (panels composed of other panels), non-disease-specific panels, and panels with less than 10 genes. The remaining gene panels (N=209) were classified into 17 disease classes using level 2 of the PanelApp classification tree. Panels without predefined classification were manually assigned to a class. We merged panels under cancer programme and tumour syndromes panels into a unique class (cancer programme+tumour syndromes). We included the gene panels classified as haematological disorders within the class haematological and immunological disorders. Final RD classes are available in Supplementary data 1-table 1.

### Gene-disease association predictions

We applied the algorithm GLOWgenes [23] to 209 selected gene panels to get a prediction of new genes associated with every RD. For clarity, we briefly describe the GLOWgenes and its functionality, although the algorithm and the benchmarking with other similar tools is fully reported in de la Fuente el at, 2022 [23]. GLOWgenes applies random walk with restart (RWWR) algorithm through more than 30 gene-gene networks using the gene panels as seeds. It first evaluates the performance of each network by splitting gene panels into training and test sets. RWWR expands the signal from the training set in every network and evaluates ability to recover genes from the test set based on efficiency and exclusivity. It classifies networks into 13 knowledge categories (regulation, coessentiality, functional similarity, phenotype similarity, etc.) to prevent redundancy. After choosing the best network performance for every category, it gives them weights and performs a new RWWR with all the genes from the gene panel. Finally, GLOWgenes integrates all rankings and weights to retrieve a gene ranking with values from 1 (higher association) to the last gene evaluated (lowest association).

From all predictions (rankings), we obtained a matrix (GLOW-matrix) with RDs (described by their gene panels) as columns and genes as rows filled with a number from 1 (best predicted association) to x (total number of genes evaluated and worst predicted association). Genes included in the input gene panel as seeds are given the value of 0. Genes lacking a predictive association with a given RD are assigned one unit greater than the lowest association value corresponding to that RD. From all predictions (rankings), we obtained a matrix (GLOW-matrix) with RDs (described by their gene panels) as columns and genes as rows filled with a number from 1 (best predicted association) to x (total number of genes evaluated and worst predicted association). Genes included in the input gene panel as seeds are given the value of 0. Genes lacking a predictive association with a given RD are assigned one unit greater than the lowest association value corresponding to that RD. GLOW-matrix is available in Supplementary data 1-table 2.

### Comparison of gene panels in terms of gene composition and their corresponding predictions

We calculated the Jaccard index between all pairs of the 209 gene panels to calculate their similarity according to their gene composition (GC-JI). To calculate the similarities between the predictions for every gene panel, Jaccard index was calculated using the N-top genes of the rank provided by GLOWgenes (P-JI), being N the number of genes in the gene panel.

We calculated the differences between the two Jaccard indexes calculated for every pair of gene panels (D-JI = GC-JI – P-JI).

### Evaluation of protein families according to the similarity in their predicted disease associations

Genes were annotated to protein families defined by the Panther resource (v19) [25]. We used Panther annotation levels 1 and 2 (Supplementary data 1-table 3), named protein families and subfamilies respectively. We performed a t-SNE analysis to the GLOW-matrix to show similarity among genes according to their rank in the predictions of association with the diseases. Genes were annotated and colored according to their protein families. In a second analysis, we performed a new t-SNE in every protein family filtering the GLOW-matrix accordingly and coloring genes as protein subfamilies.

### Clustering of genes according to their predicted association to rare diseases

We applied k-means to the GLOW-matrix to extract five clusters of genes with similar behavior in their predicted disease associations. K=5 was chosen to describe top and bottom associated genes predicted over the set of RDs and three levels of medium ranged behaviors. For visualization purposes, for each gene we calculated: 1) the percentage of diseases where they are ranked at the top of the predictions (5%); 2) the median of its position in the rankings. Values of 0 in the GLOW-matrix were not considered for these calculations as they indicate being the seeds of the predictions, that is, already known disease associations. BiomaRt 2.60.1 [26] was used to annotate the gene type of the loci with Ensembl release 114 [27]. EnrichR [28] was used for functional enrichment.

### Obtaining disease-specific and disease-class-specific genes

For this task, we used the most specific gene panels, filtering out gene sets that fully included the smallest ones, leaving a total of 183.

To identify disease-specific genes, we classified genes within each gene panel ranking as top if it fell within the top 1%, and bottom if it fell below the 50%. Genes were considered disease-specific if they were classified as top in only one gene panel and bottom in at least 90% of the remaining gene panel. To identify disease-class-specific genes, we set the top threshold to the top 5%. A disease class was classified as top or bottom if at least 50% of the diseases within that class fell into the corresponding category. Genes that ranked top 5% exclusively in one disease-class and within the bottom category in more than 10 other disease-classes, were classified as disease-class-specific.

### A gene-disease specificity score

We extracted the difference between the best ranking position (Rbest) and the median ranking position (Rmed) for each gene in the whole set of predictions. To allow comparisons across all genes, these values were normalized to the range between 0 and 1, obtaining the score of gene disease specificity (SGDS).

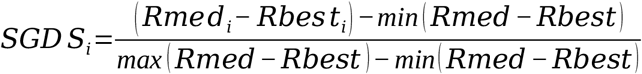.Where min(Rmed-Rbest) and max(Rmed-Rbest) correspond to the minimum and maximum differences observed across the full gene set.

A high SGDS indicates a broader distribution of ranks and, consequently, a higher specificity associated with the disease where it achieved the best ranking (Rbest).

## RESULTS

### A collection of candidate genes associated with rare diseases

A major challenge in the diagnosis of RDs is the discovery of new gene-phenotype associations. *In silico* predictions may aid in the prioritization of hypothesis to be further tested, especially when using WES or WGS. To provide a new catalog of candidate genes in RDs and be able to analyze predictions globally, we selected 209 RDs defined by gene panels from the PanelApp resource and classified them into 17 RD classes (Figure 1A). The number of RDs per class ranges from 2 in *hearing and ear disorders* and *growth disorders*, to 44 in *neurology and neurodevelopmental disorders* (Figure 1A). In total, 4,414 genes are associated with at least one of the 209 RDs, most of which are included in one or a few gene panels (Figure 1A).

**Figure 1.**
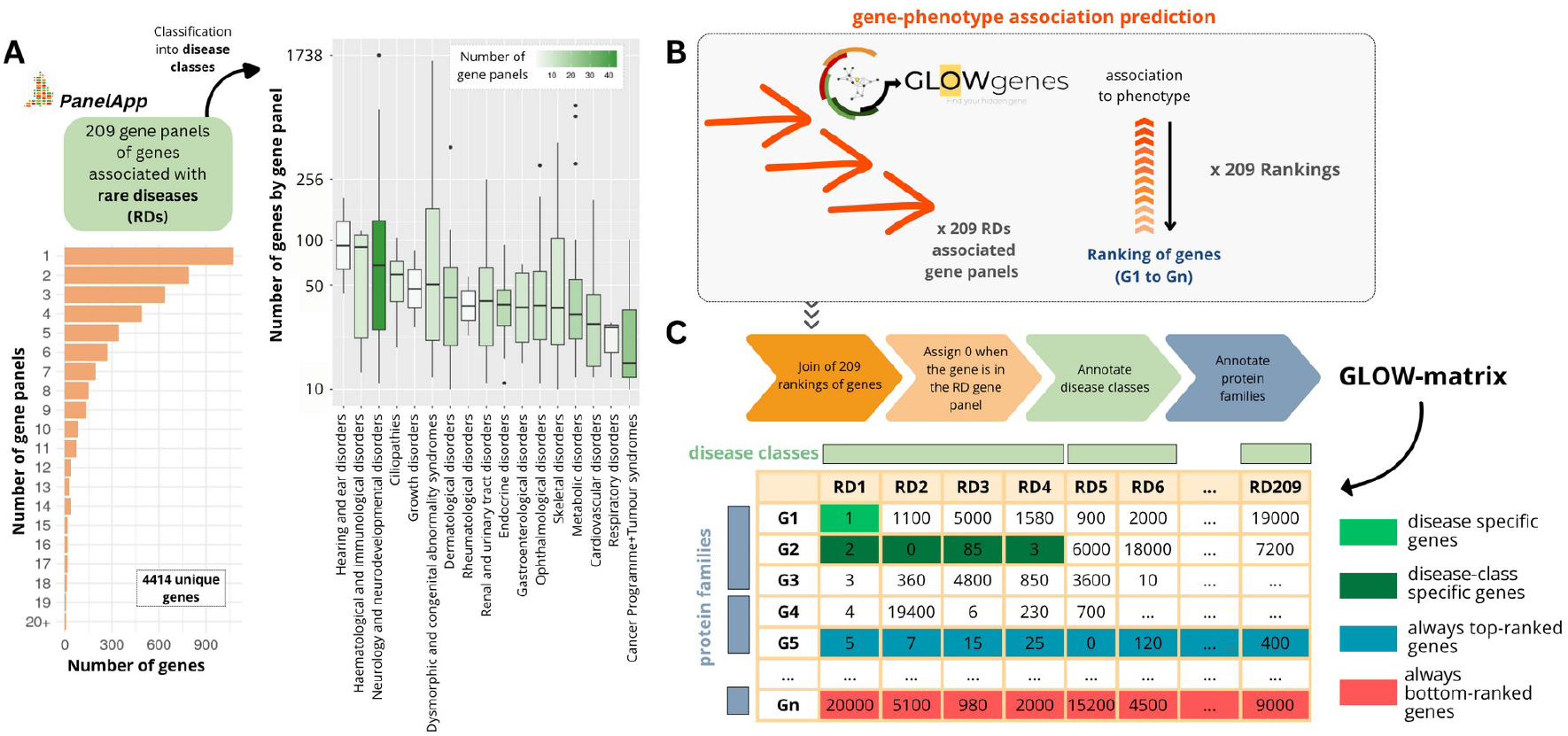
Collection, classification and analysis of rare diseases (RDs) associated gene panels. (A) Boxplot with the number of genes included in the 209 gene panels associated with RDs in the PanelApp resource. Each gene panel is classified into one of 17 RD classes. The intensity of green color indicates the number of gene panels included in each RD class. The orange bar plot shows the distribution of genes by the number of panels they are included in. (B) GLOWgenes was applied to generate a ranked list of genes predicting gene-RD associations for each of the 209 gene panels. (C) Generation of a matrix with information about RDs and genes (Gs) known and predicted associations (GLOW-matrix). Examples of disease-specific, disease-class-specific and non-specific predicted genes are shown.

We applied a Systems Biology-based algorithm, GLOWgenes, to generate new predictions of associations between genes and 209 RDs, obtaining a ranked list of genes associated with each RD (Figure 1B). Across all rankings, GLOWgenes retrieved 24,757 loci, including genes, pseudogenes and non-coding RNAs (ncRNAs). Using this data, we constructed a matrix (GLOW-matrix) with gene panels as columns and genes as rows (Figure 1C). The GLOW-matrix was populated with the ranking positions obtained from each prediction, assigning zeros for known gene-RD pairs and a value one unit higher than the lowest ranking position for gene-RD pairs not retrieved.

The GLOW-matrix represents the complete repertoire of gene-RD known and predicted associations. It is provided in Supplementary data 1-table 2. The GLOW-matrix is used to extract genes with broad associations across diseases as well as genes linked specifically with one or few diseases (Figure 1C).

### Rare diseases sharing known associated genes can exhibit divergent predictions, and vice versa

*In silico* predictions based on network propagation for gene-phenotype associations are sensitive to the set of genes used as input. While it is expected that similar gene panels yield similar predictions, we hypothesize that even small differences in the seed gene set can lead to significant divergence in the predicted genes associated with a given phenotype. In order to test this premise, we calculated two types of similarity metrics between each pair of gene panels using the Jaccard index: (1) Gene Composition Jaccard Index (GC-JI), measuring the overlap of known genes between panels, and (2) Prediction Jaccard Index (P-JI), measuring the similarity of their n-top predicted genes as performed elsewhere [29] [30]. We then computed the difference between GC-JI and P-JI (denoted D-JI) to assess how much the predictions deviate from what would be expected. A positive D-JI indicates similar gene panels with divergent predictions, while a negative D-JI reflects dissimilar panels that converge on similar predictions.

Figure 2 shows the correlation between GC-JI and P-JI. We highlight RD pairs with the most extreme differences between gene composition and prediction similarity. For instance, *ductal plate malformation* vs. *polycystic liver disease* shares a high proportion of known associated genes (GC-JI = 0.71; 15 genes in common) but show minimal overlap in predicted genes (P-JI = 0.09; 3 shared predictions). Other pairs of gene panels with similar composition but divergent predictions include: *inherited phaeochromocytoma and paraganglioma* vs. *inherited phaeochromocytoma and paraganglioma excluding NF1* (GC-JI = 0.70, P-JI = 0.21); and *early onset or syndromic epilepsy* vs. *intellectual disability* (GC-JI = 0.37, P-JI = 0.2, with 681 known and 414 predicted genes in common).

**Figure 2.**
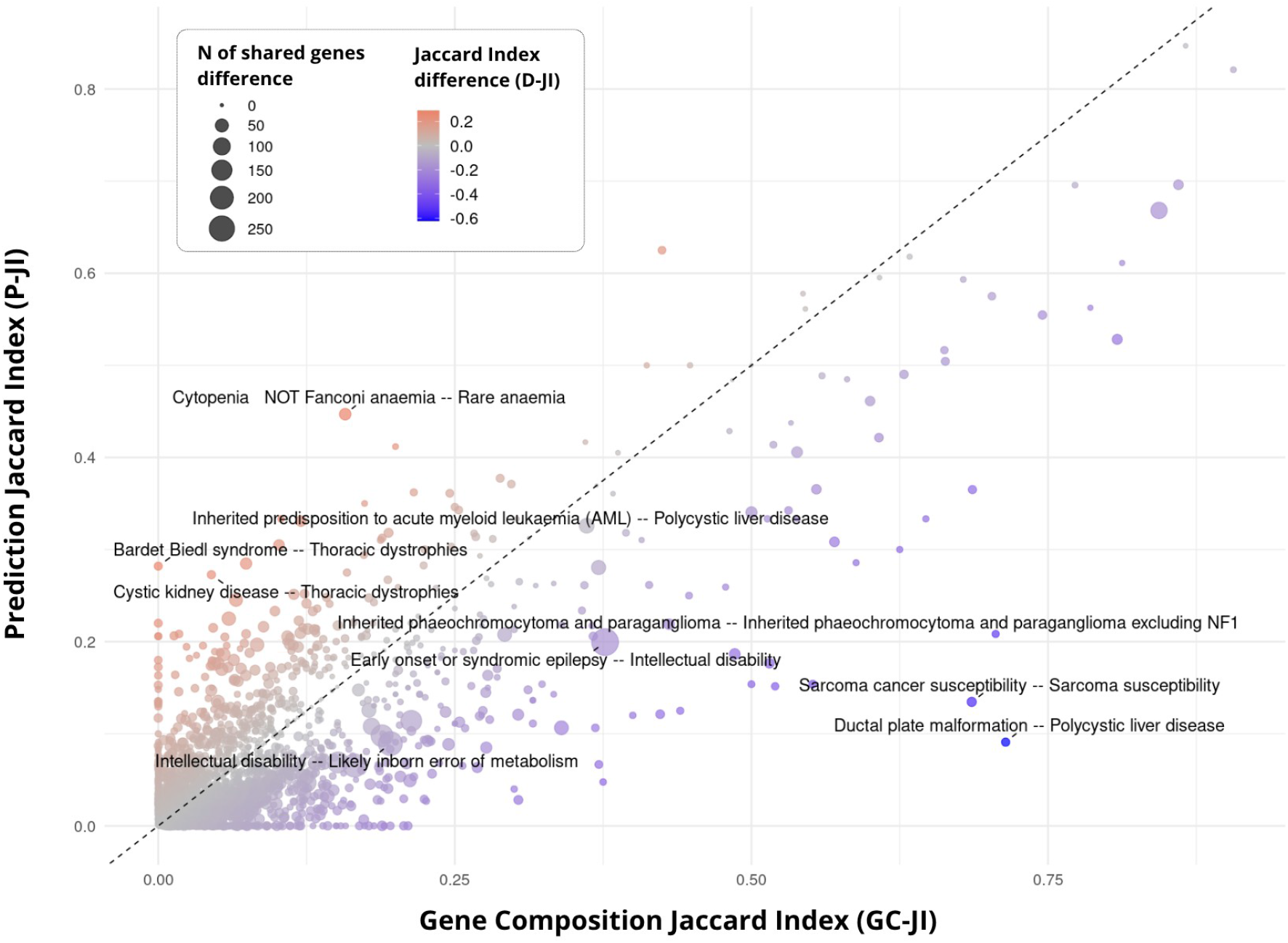
Convergent and divergent predictions. Correlation between the Gene Composition Jaccard Index (GC-JI), which quantifies the overlap of known genes between gene panels, and the Prediction Jaccard Index (P-JI), which measures the similarity of their predicted gene sets. Each point represents a pair of gene panels (rare diseases). The color gradient reflects the difference between GC-JI and P-JI (D-JI), indicating divergence or convergence between known and predicted gene sets. Point size corresponds to the difference in the number of shared known genes between the two panels.

Conversely, we also identified pairs of gene panels with little to no overlap in known genes but converging predictions. For example, *Bardet-Biedl syndrome* and *thoracic dystrophies* do not share any known associated genes, yet they share 11 predicted genes among their top-ranked predictions (P-JI = 0.28). Additional convergent examples include *cystic kidney disease* vs. *thoracic dystrophies* (GC-JI = 0.04, P-JI = 0.27) and *inherited predisposition to acute myeloid leukemia* (AML) vs. *polycystic liver disease* (GC-JI = 0.10, P-JI = 0.33).

### Some protein families also exhibit divergent predictions

Protein families are known to define functional units [27]. We wanted to explore whether these functional entities also determine behaviors in guilty-by-association based predictions, we performed a t-SNE analysis on all genes in the GLOW-matrix, capturing their similarity. Genes were annotated into Panther protein families and subfamilies [25]. Although the overall t-SNE projection reveals some patterns (Figure 3A), we further analyzed the data at the subfamily level to uncover finer-grained organization (Figure 3B). This revealed that certain protein families contain subfamilies that are clearly dissociated in the embedding space. For example, the *DNA-binding transcription factor* family comprises three subfamilies (*helix-turn-helix, immunoglobulin fold*, and *zinc finger transcription factors*) whose proteins cluster within each subfamily and show different patterns of disease association. A similar pattern is observed in *intracellular signal molecules, cytoskeletal proteins*, and *transporters*. In contrast, families like *metabolite interconversion enzymes* and *protein-modifying enzymes* display more homogeneous distributions among their subfamilies, although some degree of grouping is still present.

**Figure 3.**
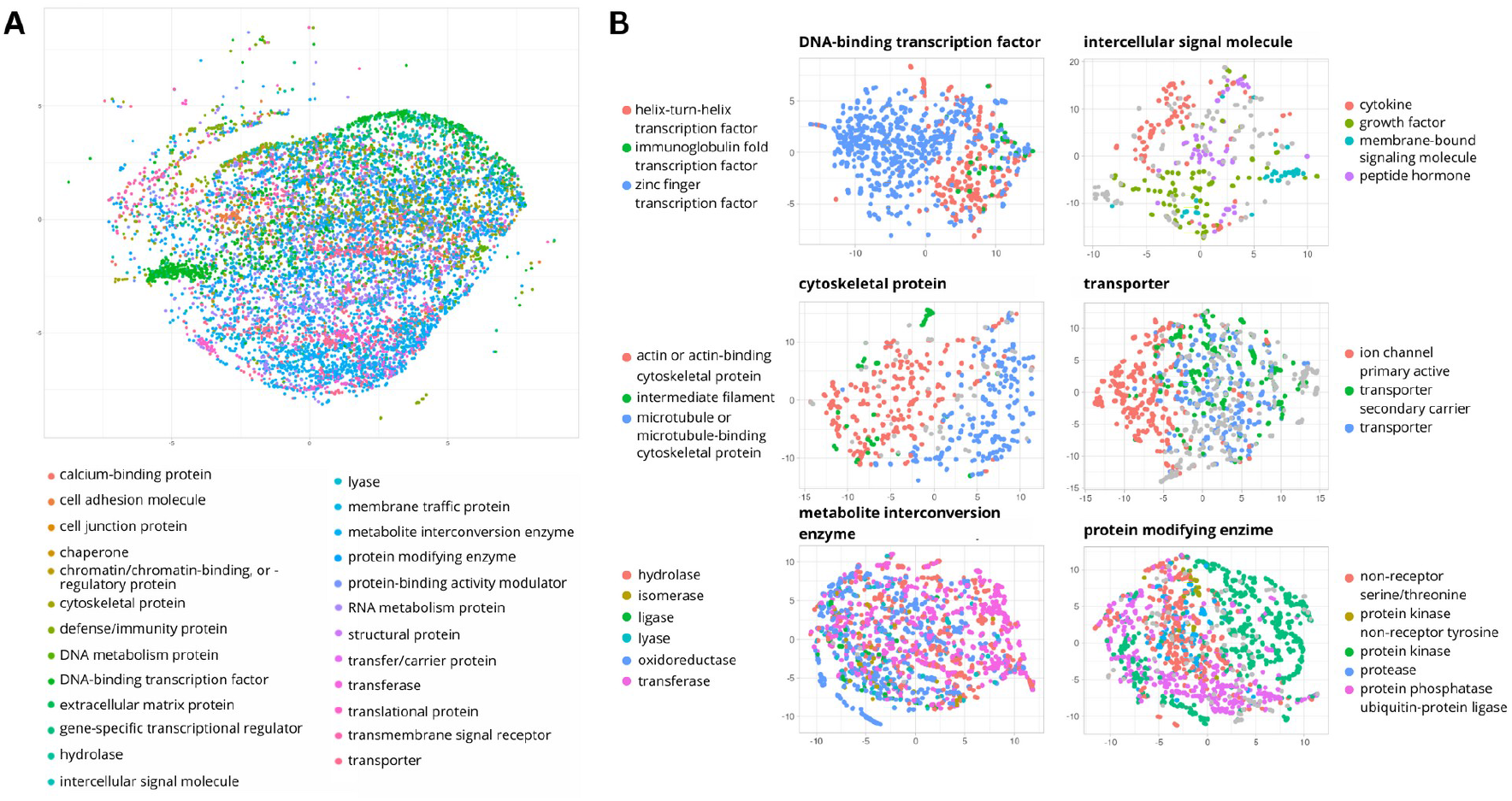
Gene-phenotype association patterns behavior within protein families. (A) t-SNE of all gene-phenotype association prediction ranking value colored by protein family. (B) t-SNE of the subset of protein family genes colored by protein subfamily.

### Genes with wide or poor predicted associations to rare diseases

Once we analyzed the behavior of gene panels and protein families in the predictions, we zoomed in to genes. To identify genes with a similar behavior in their predicted association to RDs, we applied K-means clustering to obtain five groups of genes for further analysis (Figure 4A) with RDs. Our interest focused on the top and bottom genes; therefore, we chose K = 5 as a reasonable compromise. A clustering with K=7 provided similar results (Supplementary data 2). Genes in cluster 1 are likely to appear frequently among the top predictions, regardless of the RD. In contrast, genes in cluster 5 include genes with the lowest mean ranking positions, indicating poor association scores across RDs. Clusters 2, 3 and 4 represent intermediate scenarios. Loci other than protein-coding genes are mainly found in cluster 4 and, most prominently, in cluster 5.

**Figure 4.**
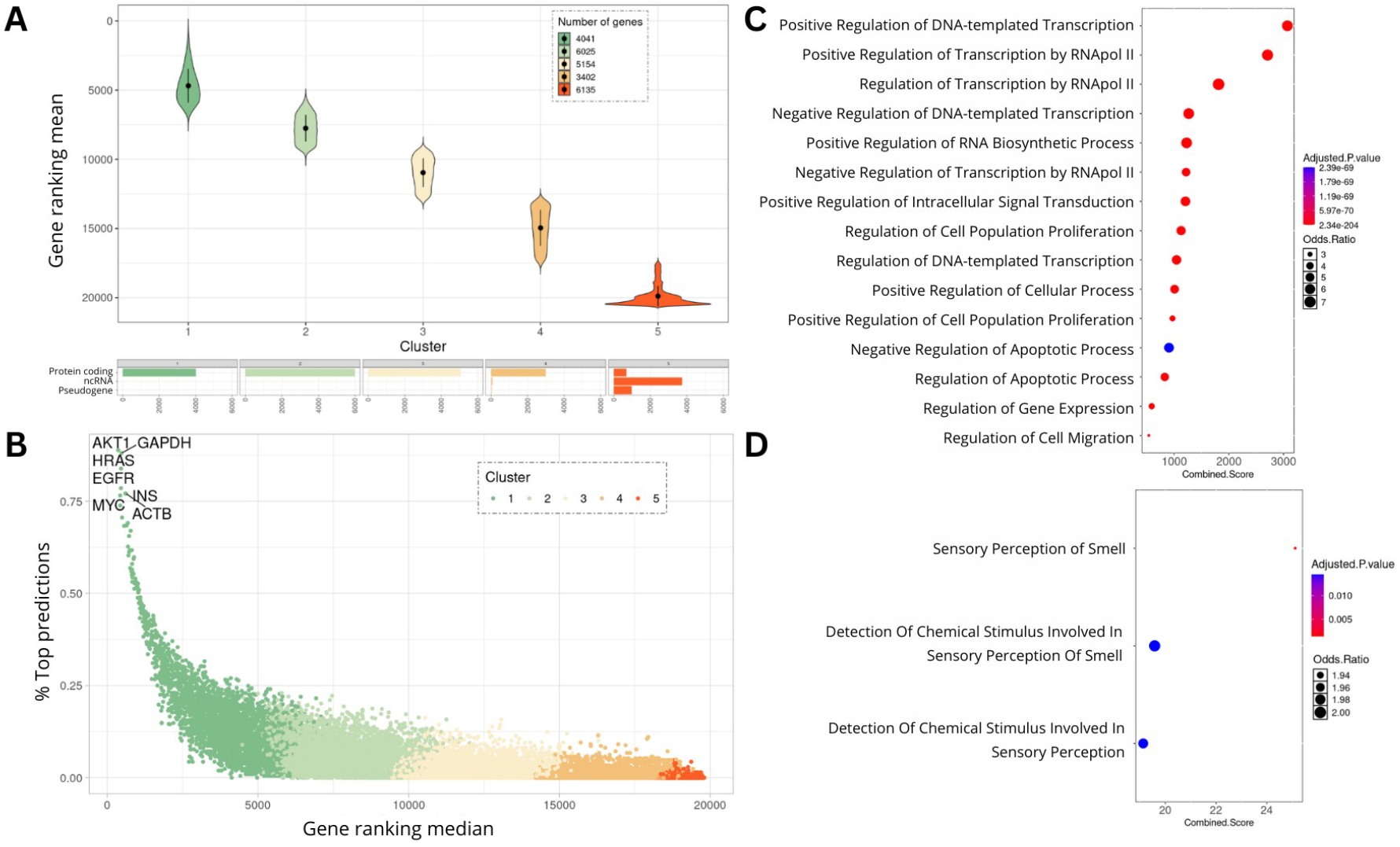
Genes and functional features systematically associated and not associated with rare diseases. (A) Boxplots with the distribution of the mean ranking values of the genes, grouped by identified clusters. Includes the composition of each cluster by loci type. (B) Scatter plot of genes ranked by the percentage of top associations across diseases (% Top Predictions) versus their overall median ranking across all rare diseases. (C) Gene ontology enrichment analysis genes in cluster 1 (most linked to diseases). (D) Gene ontology enrichment analysis genes in cluster 5 (less linked to diseases).

To extract the genes most widely predicted to be associated with RDs, we calculated, for every gene, the percentage of RDs in which it appears within the top 5% of ranked predictions. We then correlated this percentage with the gene’s median ranking position across all RDs (Figure 4B). Genes with a high median rank (i.e., closer to the top of the rankings) tend to be strongly associated with many RDs, whereas those with lower median ranks are associated with few or no diseases. The gene most consistently prioritized across RDs is *AKT1*, appearing in the top 5% of predictions for 88.8% of RDs and having a median ranking position of 362. Next highly ranked genes are *GAPDH* (88.03%, median = 469) and *HRAS* (% Top Prediction = 83.85%, median = 456). We extracted functions enriched in genes from clusters 1 and 5. Central biological functions such as positive regulation of DNA-templated transcription, regulation of apoptotic process or regulation of gene expression are enriched in cluster 1 (Figure 4C). Three functions are significantly enriched in cluster 5; none of them are key biological roles (Figure 4D).

### Disease-specific and disease-class-specific associated genes

In the molecular diagnosis of RDs, if no pathogenic variants are found at genes already associated with the patient’s phenotype, alternative variants might be considered and further evaluated. This can lead to the proposal of new candidate genes potentially linked to the RD. Computational methods play a key role in prioritizing these candidate genes. Regardless of the tool used, a valuable complementary information would be to understand how specific or broadly a gene tends to be predicted in association with RDs.

We classified a gene’s ranking position in a RD prediction as either top or bottom, being top genes those within the top 1% of the genes ranked, and bottom genes, those below the 50%. Of all the genes, 62.85% did not achieve any top positions, 12.2% achieved a top position in one RD, and 6.5% in two RDs (Figure 5A). Resuming clustering of genes according to their overall predictions, genes at cluster 4 and 5 seem to be an important source of RD specific genes (Figure 5B).

**Figure 5.**
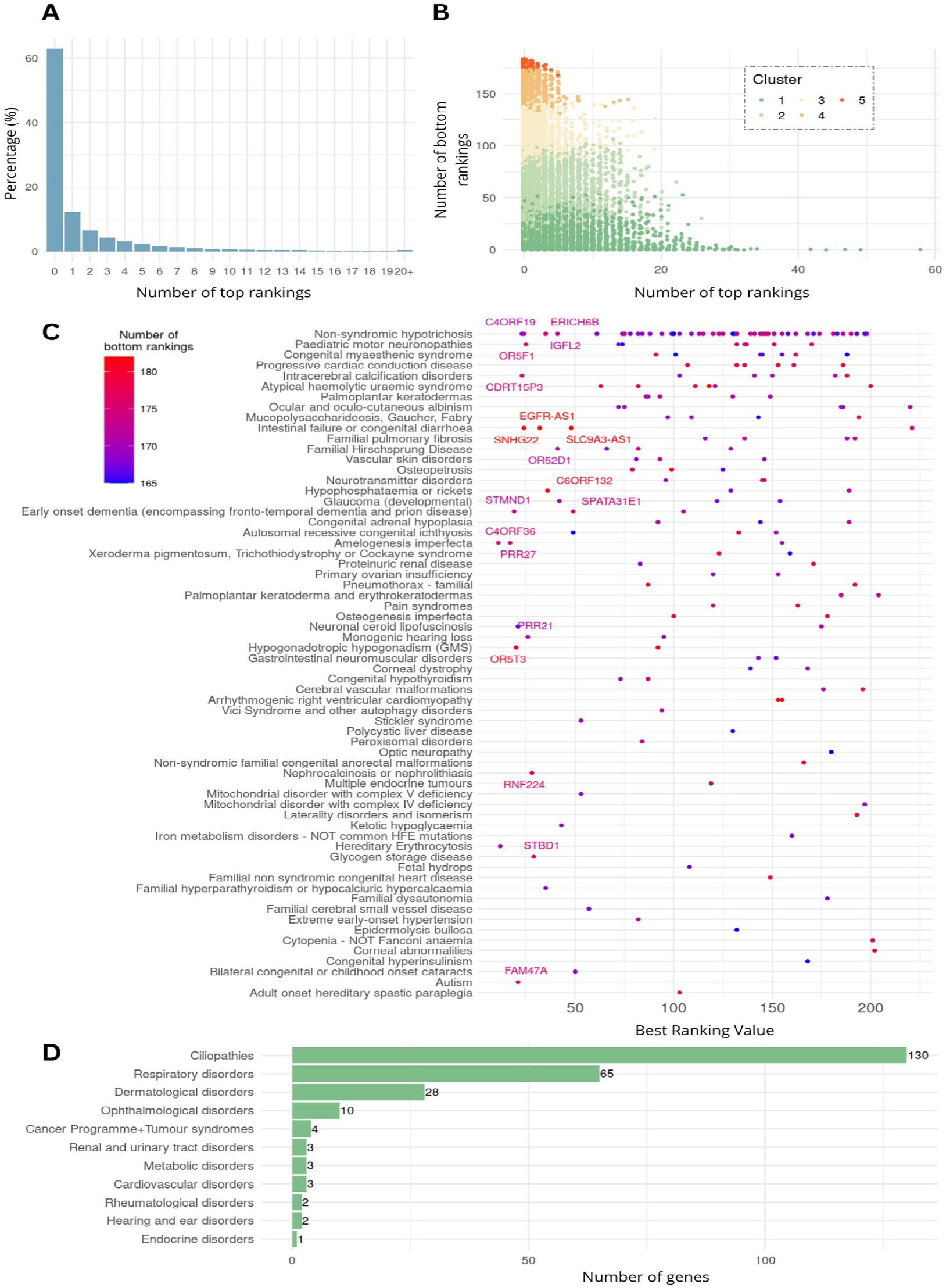
Disease-specific and disease-class-specific genes. (A) Percentage of genes classified by their number of top gene-phenotype association rankings. (B) Number of top/bottom gene-phenotype association rankings achieved by gene. (C) Description of the disease-specific genes found associated with each rare disease, plotted by their best ranking value and their number of bottom gene-phenotype association rankings. Gene names are included for those with a best ranking position < 50. (D) Number of disease-family-specific genes belonging to each family of rare diseases.

We defined disease-specific genes as those classified as top for only one RD and as bottom in 90% of the remaining RDs. Up to 192 disease-specific genes were found (Supplementary data 1-table 4). The three diseases with the highest number of disease-specific genes are: *Non-syndromic hypotrichosis* (48 genes); *paediatric motor neuronopathies* (8 genes); and *congenital myasthenic syndrome* (7 genes). We highlight disease-specific genes with extreme behaviors, a very high top position (over 50 top genes) and coloured by the decreasing number of RDs classified as bottom (Figure 5C).

We also defined disease-class-specific genes. For this task we expanded the top gene threshold to the top 5% of the ranking. A gene is considered top or bottom for a disease class if it held that position in at least 50% of the diseases within that family. Thus, we defined disease-class-specific genes as those classified as top in only one family of RDs and as bottom in more than 10 of other families of RDs. A total of 251 disease-class-specific genes were found (Supplementary data 1-table 5), 51.79% associated with ciliopathies, 25.87% with respiratory disorders, 11.15% with dermatological disorder and 3.98% to ophthalmological disorders (Figure 5C).

### A Specificity score for prioritizing potential gene–rare disease associations

If a new gene is hypothesized as a candidate for association with a RD, regardless of the methodology used to identify it, it would be informative to assess how specific this potential association is to that particular RD. Thus, we defined the Score of Gene-Disease Specificity (SGDS) as the difference between the median ranking position across all analyzed RDs (Rmed) and the best ranking position observed (Rbest). This difference is then normalized to a range between 0 and 1. A value of 0 indicates non-specific genes whose ranking positions show little variation across RDs, whether consistently ranked at the top, bottom, or mid-range. A SGDS of 1 reflects highly specific genes that reach top-ranking positions in only a few RDs, suggesting a more exclusive association. SGDS is designed to be used to sort genes with a high association score from GLOWgenes, or any other predictor, in order to highlight those that are more specific to a disease which are also presumably the less obvious (Figure 6A).

**Figure 6.**
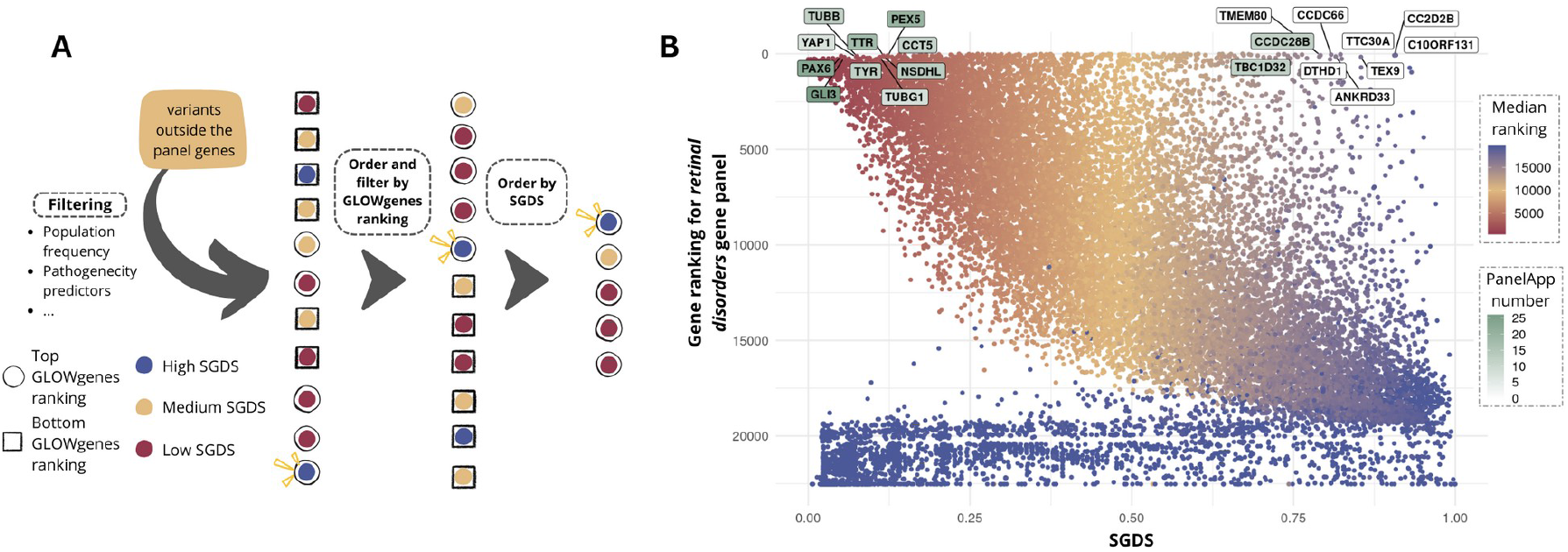
Application of the Score of Gene-Disease Specificity (SGDS). (A) Schematic workflow for gene prioritization process using GLOWgenes ranking and SGDS. (B) Application to genes predicted to be associated with retinal disorders. In the y-axis the genes are ranked as their predicted association to the disease. In the x-axis the genes are ranked as their SGDS. Point color corresponds to their median position in the rankings representing the predictions of association to >200 rare diseases. Gene names are labeled for the top 10 with the highest and lowest SGDS with a predicted association ranking < 200. The green intensity of their label corresponds to the number of gene panels from PanelApp in which the gene is included.

To evidence the utility of the SGDS, we used the genes’ ranking obtained as prediction to retinal disorders as a proof of concept (Figure 6B). We observed that some genes predicted to be associated with retinal disorders, (within the below 200 in this case first ranked genes), exhibited low SGDS due to their broader predicted association to multiple RDs (*PAX6, GLI3, YAP1, TUBB, TYR, among them*). Notably, these genes are present from 7 to 26 PanelApp gene panels (Figure 6B). In contrast, the SGDS highlights others (TMEM80, CCDC28B, TBC1D32, CCDC66, DTHD1, ANKRD33, TTC30A, TEX9, CC2D2B, C10ORF131) as disease-specific genes, which are either exclusively associated with retinal disorders or only linked to a limited number of other RDs. Among these, only *TBC1D31, CCDC66*, and *DTHD1* are included in PanelApp, appearing in 12, 12, and 1 gene panels, respectively. Additionally, *DTHD1* is listed among the red genes of the retinal disorders panel, while *TBC1D32* has been recently added to the amber genes of the retinal disorders panel in version 8.5, a release after the one used in this study.

## DISCUSSION

The application of WES and WGS as first-tier diagnostic tests for RDs introduces new challenges, ranging from data storage to results interpretation and clinical integration [31]. One of the main challenges is managing thousands, or even millions, of genetic variants, that, in the absence of known genotype-phenotype associations, are all potentially suspicious *a priori*. Given the vast number of possible hypotheses, the performance of functional assays must be strategically guided. Currently, evidence gathering and the “guilty by association” principle represent the most effective strategy for bridging the gap between data and information [32]. The main challenge in these strategies is to overcome the bias introduced by the most extensively studied genes [33]. This work aims to contribute to the genotype-phenotype discovery in RD diagnosis through a global survey and a comprehensive analysis of gene-RD associations predictions that provide us with a broader view that can enhance and accelerate diagnosis. We chose GLOWgenes as predictor, an algorithm that we know deeply, for practical reasons and mainly because it has been benchmarked against state-of-the-art tools [23], showing improved overall performance in RDs [34]. Still, results do not intend to evaluate its performance, and our conclusions and deliverables are not specific to the predictor used.

Incorporating active gene discovery capabilities into sequencing analysis pipelines may indeed contribute to new diagnosis. However, to our knowledge, only PanelApp [22] compiles candidate gene sets (called red genes) for a range of RDs that can be used to filter variants in a genetic test after discarding genes already associated with the phenotype. Tools such as Phenolyzer [21] can generate personalized gene panels for individual patients, but no current resources offer genome-wide predictions tailored to specific RDs. Our first contribution in this work is genome-wide rankings of genes based on their predicted association with 209 RDs. This dataset updates and expands our previous predictions [23] and constitutes the most comprehensive resource available for candidate genes in RDs. What remains uncertain, however, is which specific cases they may help to solve, given the rarity, disperse distribution, and, in some cases, underdiagnosis or neglect of these conditions [35].

We applied a heterogeneous set of biological networks to predict novel genes associated with each RD, as defined by PanelApp. As expected, similar gene panels tend to yield similar prediction patterns. However, the use of such an intricate network also reveals both convergent and divergent prediction behaviors across pairs of panels. At one extreme, we highlight the case of *polycystic liver disease* (PLD) and *ductal plate malformation* (DPM), two disorders originating from defects in the ductal plate [36], the embryonic precursor of the intrahepatic bile ducts. Although PanelApp defines them sharing 15 out of their 21 causal genes, they are clinically distinct, PLD manifesting later in life and often benign, and DPM with early onset and associated with more severe hepatic complications. In consonance with this clinical divergence, our predictions for these two panels overlap in only three genes. Conversely, the pair of diseases *Bardet–Biedl syndrome* (BBS) and *thoracic dystrophies* (TD) illustrate a convergent example. Both are syndromic ciliopathies that share few clinical features, such as polydactyly and renal involvement. Although, they do not have any causal genes in common according to PanelApp, they include variants in genes involved in intraflagellar transport [37][38]. On top of this, of the 20 genes listed in BBS panel, 15 of them are included into the candidate (red) genes for TD in PanelApp.

Along the same lines, we also observed protein families with subfamilies with either consistent or divergent prediction patterns. For example, enzyme classes, such as metabolite interconversion enzymes and protein-modifying enzymes, tend to define widespread, functionally broad subfamilies. In contrast, regulatory proteins (e.g., DNA-binding transcription factors), signaling proteins (e.g., intercellular signal molecule), and structural proteins (e.g., components of the cytoskeleton or transporters) tend to form more functionally specific groups. This observation suggests that “guilt by association” assumptions based on protein family membership may be informative in some cases but are not always reliable in the context of rare diseases [39].

Clustering analysis identified a subset of genes that consistently ranked highly in the prediction of associations with RDs. At the top five were AKT1, GAPDH, HRAS, EGFR and INS, genes known for their central biological roles [40][41][42][43] and extensively studied and described for their implication in cancer [44][45] and diabetes [46]. Their association to RDs is not necessarily spurious, they are extensively listed as causal and candidates in several RDs by PanelApp. At the other end of the spectrum, we identified a cluster of genes that consistently ranked among the lowest. This group is enriched in non-coding elements, which typically lack functional annotation. Nonetheless, it includes genes that, while scoring poorly overall, are strongly associated with a specific RD or a closely related group of RDs. As either poorly annotated genes or peripheric in the gene-gene functional networks, their role in diseases is harder to be highlighted by hightroughput experiments and in silico algorithms [23]. We believe these genes are of particular interest to RD research communities and should be shared and considered for inclusion in candidate gene panels.

We extracted 192 disease-specific genes. Some candidates to discuss include the Starch-Binding Domain-Containing Protein 1 (STBD1), found specific to glycogen storage diseases, a group of monogenic disorders that share a defect in glycogen synthesis or degradation [47]. STBD1 has been proposed to be involved in glycogen metabolism by binding to glycogen and anchoring it to membranes, thereby influencing its subcellular localization and its intracellular trafficking to lysosomes [48]. Another gene of interest is RIPOR3, which is predicted to be associated exclusively with monogenic hearing loss. This gene belongs to the RIPOR family [49] which includes RIPOR2, also known as FAM65B, and is essential for hearing function [50][51]. We also identified Stathmin Domain Containing 1 (STMND1) as specifically associated with Early onset dementia (encompassing fronto-temporal dementia and prion disease) gene panel. STMND1 is part of the stathmin family, small and unstructured proteins that bind tubulin dimers and are implicated in several human diseases. Although the functions of canonical stathmins (STMN1–4) remain incompletely understood and show variable expression patterns [52], STMN2 has been linked to motor neuron disease, amyotrophic lateral sclerosis [53] as well as to neurodegeneration and frontotemporal dementia [54][55].

We also report 251 disease-class-specific genes. Here we find the Ubiquinol-Cytochrome C Reductase, Complex III Subunit X (UQCR10), predicted to be associated with eight metabolic disorders. UQCR10 is listed as candidate gene in three mitochondrial-related PanelApp gene panels, all categorized under metabolic pathologies [56]. Another notable case is Zinc finger gene ZNF385C, found exclusively associated with gene panels from hearing and ear disorders, and previously proposed as candidate gene for deafness [57]. The disease class with the highest degree of gene specificity was ciliopathies. Here we report the Coiled-Coil Domain-Containing 28B gene (CCDC28B), included as a candidate (red gene) in several ciliopathies PanelApp gene panels. CCDC28B has also been described as a genetic modifier of Bardet-Biedl syndrome 1 [58].

Besides the proposal of particular new gene-RD associations, we wanted to wrap all the knowledge generated into a single score able to define disease specificity in our predictions as a proxy to detect the less obvious still strong associations. SGDS is designed to be integrated into a WES and WGS analysis pipeline to be used as variant prioritization together with GLOWgenes, or any other predictor, scores. Applied to retinal disorders, the SGDS can distinguish between broadly associated and specific disease genes. Retinal disorders are a group of rare diseases extensively studied, with diagnosis yields around 60% [59][60], substantial overlap with other diseases and a well-established and stable consortia [61]. It is expected that next genes associated to these pathologies may lie among the less obvious candidates. SGDS is able to highlight strong predicted genes but not quite studied within the rare disease community in general.

Some limitations are worth reporting. Conclusions regarding broadly associated genes may be biased by the composition of the PanelApp resource that might have overrepresented or underrepresented RD types. To mitigate this bias, we performed a manual curation of gene panels and grouped them into broader RD families. Although GLOWgenes is optimized for RDs and introduces into the prediction model gene-gene not obvious associations (e.g. genetic interactions, mouse models-based networks or drug sharing links), our analysis relies mostly on known functional annotations. Last, while in the recovery of disease-specific and disease-class-specific genes the selection of top and bottom thresholds influences which genes are retrieved, we chose values that balance the retrieval of a substantial number of genes with a focus on capturing those showing the most extreme behaviors.

## CONCLUSIONS

We provide herein a catalog of new candidate genes for >200 RDs and a comprehensive analysis of genes widely associated with RDs, exclusively associated with one RD and to a group of RDs. Besides a deeper understanding on the behavior of predictions based on functional links, these predictions integrated together into DNASeq analysis pipelines have the potential to contribute to the resolution of RD cases.

## DECLARATIONS

## Ethics approval and consent to participate

Not applicable

## Consent for publication

Not applicable

## Availability of data and materials

All data generated by this work is included into the supplementary material.

## Acknowledgments

This work was supported by the Instituto de Salud Carlos III (ISCIII) of the Spanish Ministry of Health (PI22/00579), co-funded by European Regional Development Fund (FEDER funds) “A way to make Europe”, IMP/00019 and PMP24/00024 programs. YB is supported by a platform technician contract of ISCIII (CA24/00004), GUR is supported by a ISCIII predoctoral contract (FI23/00082) and complemented by a Fundación Humanismo y Ciencia fellowship. PM is supported by a Miguel Servet program contract from ISCIII (CPII21/00015).

## Author contributions

Study concept and design: PM and YB. Data analysis and interpretation: YB, GUR and PM. Drafting of the manuscript: YB and PM. Manuscript reviewing and editing: YB, GUR and PM. All authors have read and agreed to the published version of the manuscript.

