## Supplementary Figure 1 for "A global survey of System Biology-based predictions of gene-rare disease associations to enhance new diagnoses"

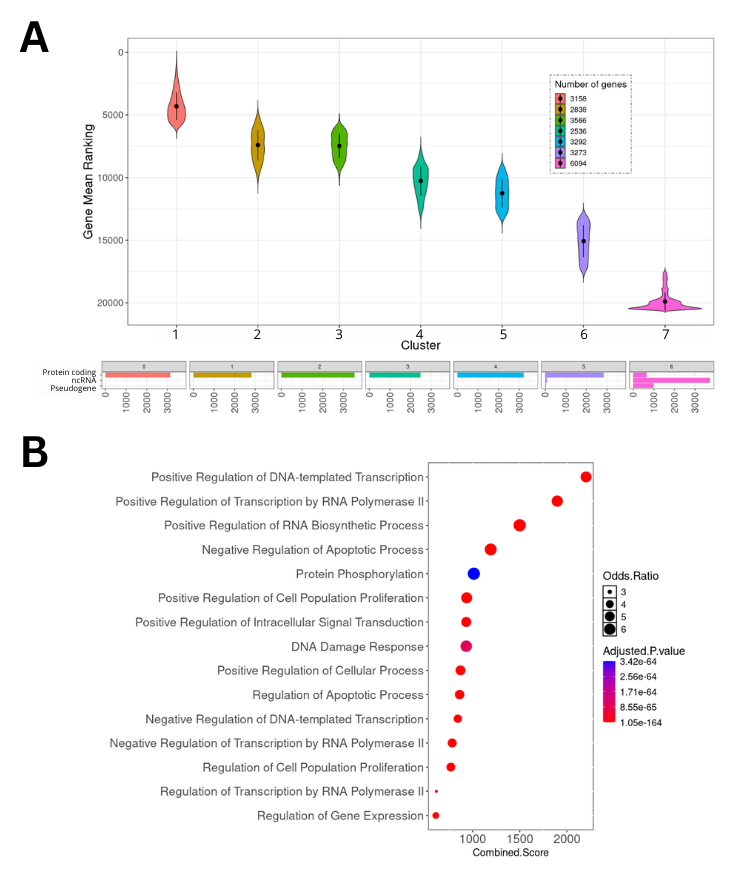

**Supplementary Figure 1. Genes and functions systematically associated and not associated with rare diseases with K=7.** (A) Boxplots with the distribution of the mean ranking values of the genes, grouped by identified clusters. Includes the composition of each cluster by loci type. (B) Gene ontology enrichment analysis genes in cluster 1 (most linked to diseases). No function was significantly enriched in cluster 7 (less linked to diseases).
